# Decoupled wing evolution reveals trait-specific links between morphology, geography, and diversification in Ithomiini butterflies

**DOI:** 10.64898/2026.09.25.754348

**Authors:** Nicolas Chazot, Maël Doré, Adrien Mathou, Keith R. Willmott, Suzie Lagarde, André V. L. Freitas, Raphael Cornette, Marianne Elias

## Abstract

Understanding when species diversification and phenotypic evolution are coupled remains a central challenge in macroevolution. Such coupling may depend on the traits considered, phenotypic integration, or the geographic context in which diversification occurs. Here, we investigate these relationships in Ithomiini, a diverse Neotropical butterfly tribe whose diversification is strongly associated with the Andes-Amazonia system. We quantified forewing and hindwing size and shape for 1,618 individuals representing 291 species using geometric morphometrics, and combined these data with phylogenetic comparative methods, diversification-rate estimates, and spatial analyses of species distributions. Evolutionary rate shifts were largely trait- and wing-specific, indicating limited coupling between forewing and hindwing evolution. Across the phylogeny, wing size evolution and extant wing size were not associated with species diversification, nor was hindwing shape evolution. By contrast, forewing shape evolutionary rates showed a positive association with diversification. Spatial analyses revealed that this coupling was geographically concentrated in the Andes, where high diversification rates coincided with elevated forewing size and shape diversity and increased forewing evolutionary rates. Hindwing diversity, by contrast, more closely tracked phylogenetic diversity and showed weaker or opposing relationships with diversification. Our results show that diversification and phenotypic evolution in Ithomiini are neither generally coupled nor fully independent, but linked in trait-specific and geographically structured ways. More broadly, they highlight the need to decompose complex phenotypes and integrate spatial context when testing macroevolutionary coupling.

## Introduction

A major goal of macroevolutionary biology is to understand whether different dimensions of biological diversification follow shared or independent evolutionary dynamics. Species diversification, phenotypic evolution, ecological differentiation, and geographic structure are often expected to interact, for instance if speciation is driven by divergent selection traits and different regions incur contrasting selection regime. Yet empirical studies show that these dynamics can be temporally lagged, spatially scale-dependent, or partially decoupled, and hence their relationships are difficult to predict (Hähn et al. 2025, Morinaga & Bergmann 2023, Chazot et al. 2021, Rabosky & Adams 2012, Folk et al. 2019, Graham & Fine 2008). In some cases, diversification and phenotypic evolution do appear to be coupled. For example, across ray-finned fishes, speciation rates and rates of body-size evolution are strongly correlated, suggesting that lineage accumulation and phenotypic change can be linked during large-scale radiations (Rabosky 2013). In contrast, a comparative study across frogs showed that diversification and phenotypic evolution are not consistently coupled, with clades spanning different combinations of lineage accumulation and morphological evolutionary rates rather than following a single macroevolutionary trajectory (Morinaga & Bergmann, 2023). These contrasting results suggest that coupling between species diversification and phenotypic evolution might be clade- or trait-specific rather than a general expectation.

Evolutionary coupling can occur not only through time, along phylogenetic branches, but also across geographic space. If species diversification and phenotypic evolution are governed by shared processes, then regions containing rapidly diversifying lineages may also contain species with elevated rates of morphological evolution or high morphological diversity. Recent work on South American freshwater fishes provides a clear example of geographically structured coupling: across the continental fauna, recent speciation rates were highest in upland lineages experiencing accelerated body-size evolution, whereas this association was not a general predictor across all environments and other drivers such as temperature, area, and diversity-dependence had limited explanatory power (Cerezer et al., 2023). Conversely, spatial mismatches between diversification rates, phylogenetic diversity, and morphological diversity may indicate that lineage accumulation and phenotypic divergence are structured by different processes.

Coupling may also occur among different structures of the phenotype. Morphological traits vary in their functional roles, developmental architecture and evolutionary lability, so different traits may show either coordinated or independent evolutionary dynamics (Klingenberg 2008, Revell et al. 2008). Accounting for multiple modalities when quantifying complex phenotypes is especially important. Size and shape, for example, may share some developmental and functional determinants, but they can also respond to different selective pressures. Body or organ size often reflects broad ecological, physiological, or life-history trait variation, whereas shape can more directly influence mechanical performance, habitat use, or behavioural function (Peters 1983, LaBarbera 1989, Betts & Wootton 1988, Le Roy et al., 2019, Vanhooydonck & Van Damme 1999, Fulton et al., 2005). Within biological structures as well, phenotypes are often organized into modules, i. e., sets of components that covary strongly internally but are relatively independent from other components (Klingenberg 2008). Morphological integration can promote coordinated evolution among traits, whereas modularity can facilitate mosaic evolution by allowing different structures to respond semi-independently to selection or constraint (Klingenberg 2008). In mantis shrimp, for example, modularity of the power-amplified raptorial appendage is associated with heterogeneous rates of morphological evolution among components, illustrating how complex functional systems can diversify through partly independent trait dynamics (Claverie et al. 2013). In this context, testing whether different anatomical components evolve together is essential for understanding how complex phenotypes diversify.

Butterfly wings provide a clear example to examine this question. Forewings and hindwings form a mechanically coupled flight apparatus but they may not contribute equally to flight performance or evolve under the same constraints. Experimental work in Lepidoptera shows that forewings and hindwings flap synchronously and are mechanically coupled, but also that flight is driven primarily by the forewing. Individuals can still fly after hindwing removal, although hindwings are important for normal evasive flight (Jantzen & Eisner 2008). Comparative analyses further suggest that butterfly wing shape is associated with flight behaviour, habitat, predators, and sex-specific behaviours, although the selective pressures underlying macroevolutionary wing-shape variation often remain difficult to identify (Betts & Wootton 1988, Le Roy et al. 2019). Recent phylogenetic work on *Papilio* butterflies also found independent evolution of forewing and hindwing shape and size, with forewings inferred to be more functionally constrained and hindwings more evolutionarily labile (Owens et al 2020). These findings make butterfly wings an especially useful system for testing whether functionally related structures show coupled or decoupled evolutionary dynamics.

Here, we investigated diversification dynamics of species and phenotypes in space and time in one of the largest and best sampled clades of Neotropical butterflies. The tribe Ithomiini comprises about 400 described species, found across South and Central America (Brown, 1979, Chazot et al., 2019, Doré et al., 2022). Diversification, however, occurred primarily in the Andes-Amazonia system (Chazot et al. 2019) and current species richness is primarily concentrated along this axis (Chazot et al. 2016, Doré et al., 2022). Ithomiini are chemically defended (Brown 1985) and selection on wing colors has led to color pattern convergence that advertises their unpalatability to predators (Müllerian mimicry, Müller 1879), both within Ithomiini and with other butterfly lineages (Beccaloni 1997, Brown & Benson 1974, Pérochon et al., 2025).

A rich scientific literature on Ithomiini and relatively comprehensive museum collections are available. These resources provide comprehensive information on taxonomy, phylogeny (Chazot et al., 2019), species distribution (Doré et al. 2022), and sources of specimens for phenotyping, making Ithomiini a rare opportunity to investigate macroevolutionary dynamics of a large group of tropical insects. Combined with a long history of diversification across broad environmental gradients, especially in the Andes, Ithomiini are a powerful system for testing how phenotypic evolution, geographic structure, and diversification dynamics are linked. By quantifying forewing and hindwing shape variation across almost 300 Ithomiini species, we specifically address the following questions:

1. Are wing size and shape evolution coupled?
2. Do forewings and hindwings evolve similarly?
3. Are phenotypic evolutionary rates associated with diversification rates?
4. Are regions of high diversification also hotspots of morphological evolution?

## Material and Methods

### Data acquisition

The dataset comprised 291 species included in the phylogenetic tree of Ithomiini (*ca.* 72% of the total diversity), with on average 5.5 individuals per species. Dorsal and ventral photographs of specimens were taken at the Museum National d’Histoire Naturelle (Paris, France) and at the Natural History Museum (London, UK). To obtain high-quality standard photographs for digitization, the sampled specimens were placed on a vertical, square, opaque acrylic sheet measuring 355 × 355 × 3 mm. The pin on the butterfly was inserted into a small block of foam glued to one side of the sheet. Digital images of each specimen for hind and forewings were taken in RAW format (5184 × 3456 pixels) using a NikonD90 camera and converted to TIFF format.

All TIFF images were processed using `tpsDig2` version 2.32 software (Rohlf, 2010). We digitized forewing and hindwing contours of 1618 individuals using a combination of 2 landmarks and 100 sliding landmarks per wing. The semi-landmarks were placed along the contour of the right forewings and hindwings using `tpsDig2`. These semi-landmarks were positioned clockwise, starting at the front end of the wing (Supplementary Figure S1).

### Procrustes alignment and data compilation

Statistical analyses were performed using RStudio software (R Core Team, 2019) using the package `geomorph` (Adams et al., 2021). The datasets corresponding to the specimens studied for the forewings (FW) and hindwing (HW) were converted into three-dimensional tables (i.e. arrayspecs matrix: landmark matrices × number of landmarks × dimensions) (`geomorph`, Adams et al., 2021). Landmarks were then aligned and overlaid using a generalized Procrustes analysis with the function *gpagen* (Dryden & Mardia, 1993 ; Adams et al., 2021) (Supplementary Figure S2). In order to reduce the dimensionality and evolutionary covariation of the data, the shape coordinates were synthesized using phylogenetic principal component analysis (pPCA) using *gm.prcomp* (`geomorph` R package, Adams et al., 2021). For both the forewing and the hindwings, we retained the first five principal components for analyses, which together accounted for more than 90% of the shape variation. We recovered wing centroid size from the Procrustes analysis that we used for wing size estimates after log transformation.

### Rate of morphological evolution through time

We estimated branch-specific rate of evolution for both wing size and shape and both wings using the R package `RRphylo` (Castiglione et al. 2018). We used *search.shift* to identify clade shifts in rates of evolution. We divided time into 1 My time bins to calculate the average rate of evolution through time. For each branch present in a time-bin, we multiplied the absolute branch rate of evolution by the length of the branch overlapping the time-bin and calculated the mean rate of evolution across branches for each time-bin. We calculated the rate of evolution through time for each shifting clade separately from the background dynamics.

### Correlation between rate of diversification and rate of phenotypic evolution

We tested for coupling between the branch-specific rates of wing size evolution, wing shape evolution and diversification. For wing size and shape, we used the results of RRphylo analyses. For diversification rate, we used `julia`’s implementation of the cladogenetic diversification rate shift model (ClaDS, Maliet et al. 2019). To test for correlations between these different evolutionary rates, we used the Cor-STRATES framework (Cooney & Thomas 2021). Cor-STRATES compares an observed correlation to a distribution of correlations obtained with simulated trait evolution.

Togenerate simulated wing size rate of evolution, we estimated phylogenetic signal with Pagel’s lambda using *phylosig* function and used it to rescale the tree (`phytools` R package, Revell 2024). We simulated wing size data on the rescaled tree using *fastBM* (`phytools` R package, Revell 2024) and estimated again branch-specific rate of evolution with RRphylo. For generating simulated wing shape rate of evolution, we estimated phylogenetic signal with Pagel’s lambda using *mvgls* (`mvMORPH` R package, Clavel et al. 2015) function and used it to rescale the tree. We simulated wing shape data on the rescaled tree using *mvSIM* (`mvMORPH` R package, Clavel et al. 2015) and estimated again branch-specific rates of evolution with `RRphylo`. We generated 200 simulated datasets that were used to calculate simulated correlations and compute two-tailed p-values for the observed correlation coefficients. We used Spearman’s rank correlation for estimating the association between rates. We used log-transformed absolute rates of evolution for all analyses. We applied the Cor- STRATES framework to test the associations between: FW and HW size rate, FW and HW shape rate, FW size rate and FW shape rate, HW size rate and HW shape rate, FW size rate and ClaDS rate, HW size rate and ClaDS, FW shape rate and ClaDS, HW shape rate and ClaDS.

### Wing size and rate of diversification

We also tested for a correlation between wing size of extant species and diversification rate. To do so, we calculated the average root-to-tip rate of diversification from ClaDS results, by averaging the branch rates along its root-to-tip path, weighting each branch by its time duration. To test for a correlation between wing size and average diversification rate, we performed phylogenetic linear regressions with the R package *phylolm* (Tung Ho & Ané 2014).

### Geographic distribution

We retrieved continuous species distribution maps for each ithomiine species from Doré et al. (2022). These quarter-degree resolution maps were generated from an initial data set of 28,986 georeferenced occurrences covering all species. Species distributions were modeled under an Ensemble framework encompassing three species distribution modeling algorithms (i.e., Random Forest, Gradient Boosting Trees, and Artificial Neural Networks), employing bioclimatic variables, elevation, and forest cover as predictors. We binarized the continuous maps into presence-absence using a ‘probability ranking’ rule (d’Amen et al., 2015) that retains only the N highest scoring species in each grid cell, N being the expected local richness computed as the sum of each local species’ probability of presence. These binary range maps were used to compute all downstream spatial analyses based on local species presence/absence in each grid cell.

### Mapping size and shape in space

We retrieved mean centroid size and mean pPC1 scores (i.e., shape) for all species and mapped the average values recorded among local species found in each grid cell of quarter-degree resolution maps. Similarly, we mapped the distribution of phylogenetic and morphological diversity in space using the Rao’s Q entropy framework (Rao, 1982). With binary presence/absence data, phylogenetic diversity represents the average phylogenetic distance between local species. For size and shape diversity, Rao’s Q entropy represents, respectively, the average centroid size difference, and the average shape distance in the morphospace, computed among local species found in grid cells.

Finally, we retrieved the root-to-tip diversification rates from ClaDS analyses, and root-to-tip evolutionary rates for size and shape from RRPhylo for each species, and mapped average rates across all grid cells. All size and shape metrics were computed for both forewings (FW) and hindwings (HW).

### Testing for spatial correlations

We tested for spatial correlations between the distribution of three blocks of variables. The first block investigated correlations between elevation, mean size, and mean shape quantified as PC1 scores. The second block focused on spatial correlations between phylogenetic and morphological (i.e. wing size and shape) diversity. The third blocks studied correlations across diversification rates and size/shape evolutionary root-to-tip rates. We performed Spearman’s rank correlations across all quarter-degree grid cells within the range of ithomiine butterflies. To account for positive spatial autocorrelation in each metric distribution, we applied Clifford’s correction (Clifford, 1989; Dutilleuil et al., 1993) to adjust effective sample size and associated degrees of freedom. Because the degrees of spatial autocorrelations detected did not lead to lowered effective sample sizes compared to true sample sizes (i.e., number of grid cells within ithomiine range), we retained initial degrees of freedom in the statistical tests.

### Effect of altitudinal niches on wing size and shape

Since we reported significant correlations between elevation and mean size/shape observed throughout ithomiine communities, we also tested for an effect of species altitudinal niche on species size and shape in a phylogenetic comparative framework. For size, we ran Phylogenetic Generalized Linear Regression (PGLS) with the R package `nlme` v3.1-168 (Pinheiro & Bates, 2025), using log-transformed centroid size as response variables. For shape, we ran a multivariate version of PGLS with the R package `mvMORPH` (Clavel et al., 2015) using the five pPC shape axes as response variables. All models included a Pagel’s lambda (λ) parameter, optimized during model fit, to account for the degree of phylogenetic signal in the data. Models were performed for both forewings (FW) and hindwings (HW).

### Data and Code availability

All scripts and files used to carry out analyses are available in the associated Zenodo archive (https://doi.org/XX.XXXX/zenodo.XXXXXXX). In parallel, scripts and files are also available on GitHub for all spatial analyses (https://github.com/MaelDore/ithomiini_morphometrics; Release v1.0.0), and for all time-related evolutionary analyses (https://github.com/XXXXXXXX; Release v1.0.0).

## Results

### Morphological variation

The first pPCA axis describing forewing shape variation showed variation from triangular, pointy shapes, elongated along the front end of the wing for positive pPCA scores to rounded wings, elongated distal margin, for negative pPCA scores (Figure 1, Supplementary Figure S3). Hindwing variation across pPC1 ranged from round wings to angular wings exhibiting a broken front edge Supplementary Figure S4. pPCA analyses showed that subtribes segregated to a much greater extent based on forewing shape variation than on hindwing shape variation (Supplementary Figure S3, S4). This FW segregation was particularly strong between Ithomiina on one side and Napeogenina, Mechanitina and Oleriina on the other. Dircennina and Godyridina, the two most species-rich subtribes, occupied a large fraction of the FW and HW morphospace. Based on Pagel’s lambda, phylogenetic signal was stronger for wing shape than wing size and in both cases stronger for the hindwing than the forewing (λ shape forewing=0.78, λ shape hindwing =0.84, λ size forewing=0.45, λ size hindwing=0.68).

**Figure 1.**
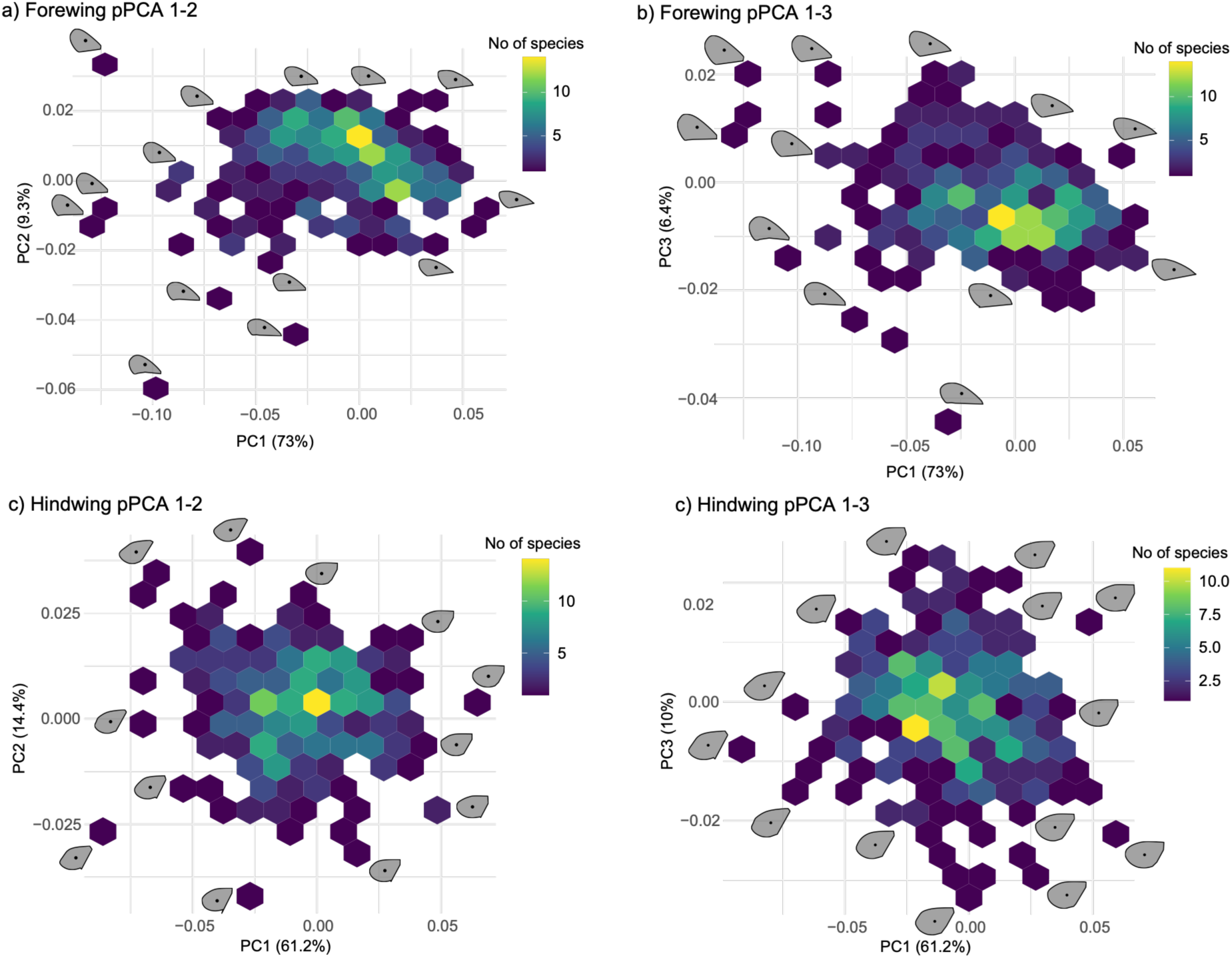
Fore and hind wing morphospace (pPCA axes 1, 2, 3), with the number of species occupying the morphospace.

### Morphological evolution through time

Using RRphylo to identify shifts in evolutionary rate dynamics of wing size evolution, we found three and two shifts for forewing and hindwing respectively, with only one shift shared between the two wings (Figure 2). We found a drop down in the rate of size evolution happening at the root of the genus *Oleria* for both the forewing and the hindwing (Figure 3). We detected two other drop downs in the rate of size evolution: at the root of the genus *Pteronymia* for forewing size and at the root of Napeogenina for hindwing size (Figure 3). We found a single increase in the rate of size evolution, for forewing, at the root of the Godyridina (*Velamysta* and *Veladyris* excluded). Wing shape showed less significant shifts of evolutionary dynamics but we identified one shift for each wing (Figures 2-3).

**Figure 2.**
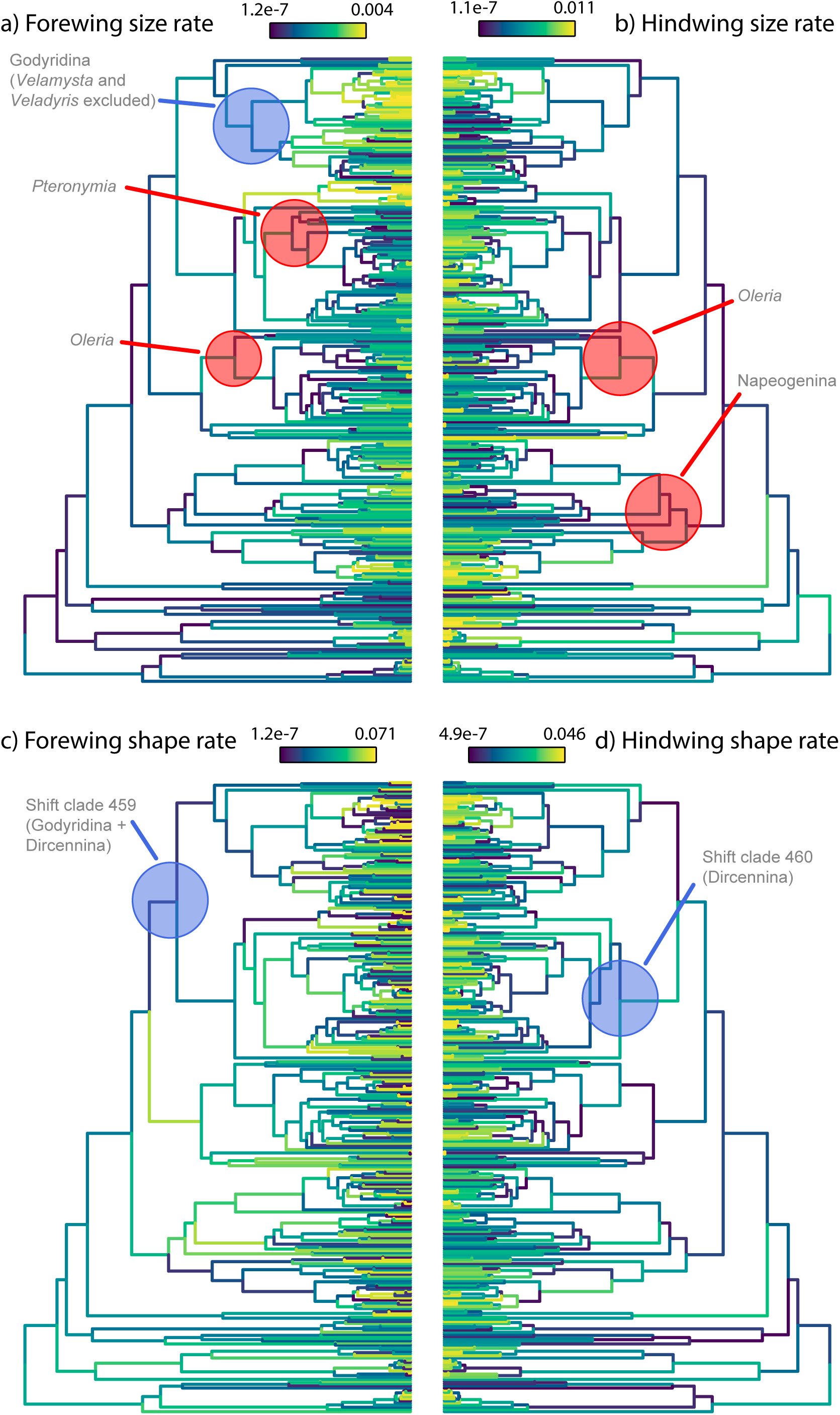
Absolute rate of evolution as estimated from RRphylo analyses for wing size (top row) and wing shape (bottom row). Shifts identified are indicated by colored circles (red=lower rate, blue=higher rate).

**Figure 3.**
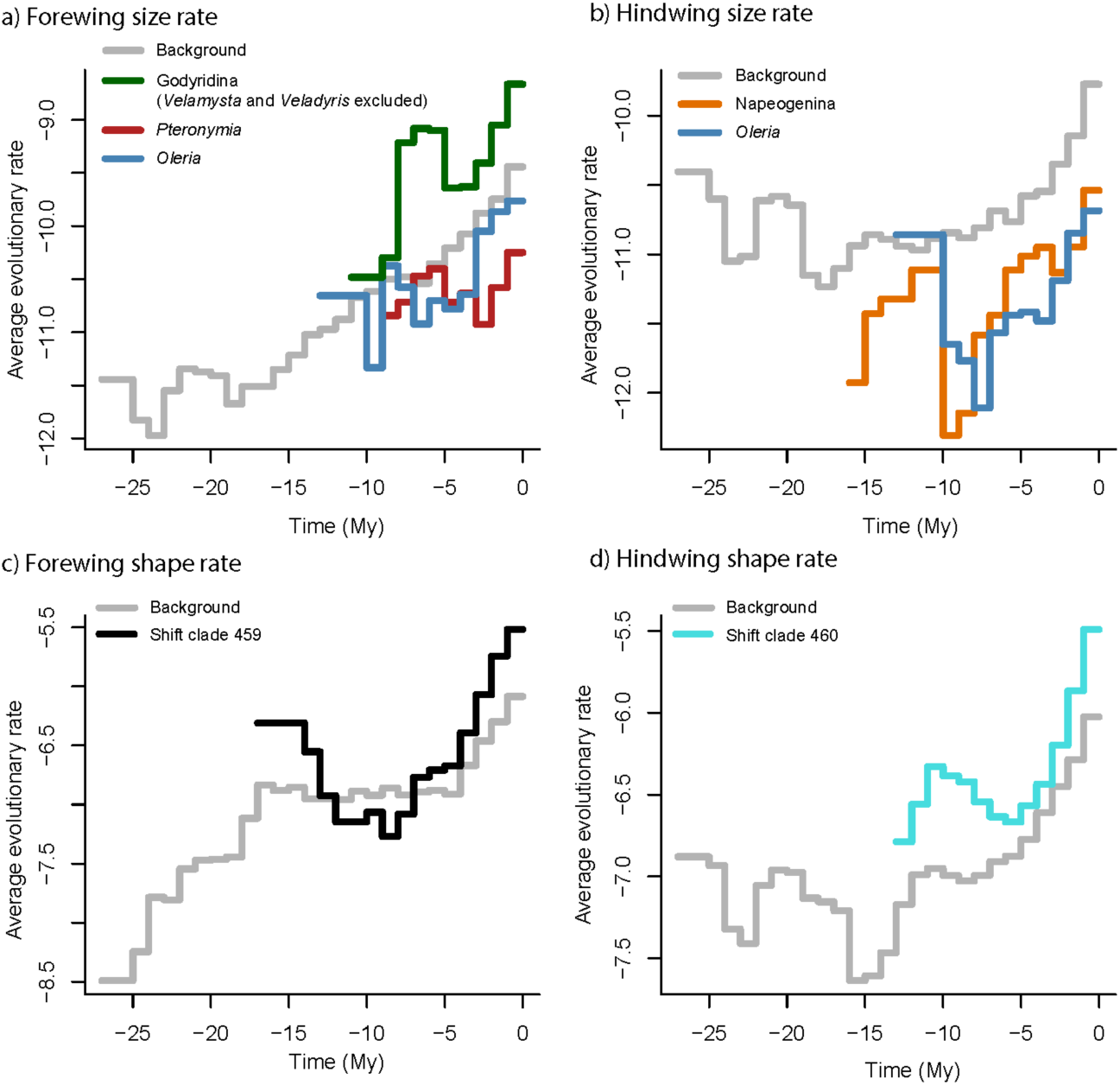
Mean absolute rate (log-transformed) of evolution through time (1 My time bins) calculated from RRphylo analyses for wing size (top row) and wing shape (bottom row). Grey lines are for the mean background rate of evolution. Colored lines correspond to the mean evolutionary rate calculated for each shifting clade.

Rate of shape evolution of the forewing increased for the clade containing the two most species rich subtribes Godyridina and Dircennina. Rate of hindwing shape evolution increased for the subtribe Dircennina only.

When measured from branch specific rate of evolution using the Cor-STRATES framework, we found diverging trends between the forewing and the hindwing, and between size and shape evolution (Table 1, Supplementary Figure S6). While forewing and hindwing shape rates showed strong correlation, this relationship was not significantly different from that expected from the phylogenetic structure only (Table 1, Supplementary Figure S6). The rates of forewing and hindwing size evolution rates were also positively correlated but Cor-STRATES analyses showed that this relationship was weaker than expected under the assumption of phylogenetic signal alone, indicating some decoupling in size evolution rates between wings. We also found positive relationships between the rates of wing size and shape evolution for each wing, but the correlation was statistically significant for the forewing only (Table 1, Supplementary Figure S6). The rate of wing size evolution showed no significant relationship with the rate of species diversification, nor did the hind wing rate of shape evolution (Table 1, Supplementary Figure S6). However, we detected a positive, albeit weak, correlation between the forewing rate of evolution and species diversification rate statistically, which was stronger than expected from phylogenetic structure alone (Table 1, Supplementary Figure S6).

**Table 1.** Correlation between evolutionary rates estimated using the Cor-STRATES framework. FW=forewing, HW=hindwing, ClaDS=rate of speciation (from ClaDS analyses), ρ_obs=observed correlation, ρ_null=mean correlation from simulated trait evolution, P=probability that the observed correlation is less extreme than simulated correlations, SES = standardized effect size of the observed correlation relative to the null distribution generated by Cor-STRATES.

| Wing(s) | Variables | $\rho_{\text{obs}}$ | $\rho_{\text{null}}$ | P (two tailed) | SES |
| --- | --- | --- | --- | --- | --- |
| HW vs. FW | Size rate | 0.358 | 0.515 | 1 | -4.451 |
| HW vs. FW | Shape rate | 0.766 | 0.764 | 0.490 | 0.081 |
| FW vs. FW | Size rate vs. Shape rate | 0.745 | 0.632 | <0.001 | 3.852 |
| HW vs. HW | Size rate vs. Shape rate | 0.654 | 0.618 | 0.098 | 1.336 |
| FW | Size vs. ClaDS | 0.061 | 0.034 | 0.228 | 0.730 |
| HW | Size vs. ClaDS | -0.017 | 0.036 | 0.776 | -1.327 |
| FW | Shape vs. ClaDS | 0.113 | 0.051 | 0.010 | 2.281 |
| HW | Shape vs. ClaDS | 0.026 | 0.051 | 0.836 | -0.931 |

Finally, we tested for a relationship between extant species wing size and diversification rate but we found no significant correlation (Supplementary Table S1).

### Morphological evolution in space

The geographic distribution of wing size and shapes revealed highly contrasting patterns between the forewing and the hindwing (Figure 4). Large forewings were clearly associated with Andean and Central American distribution whereas the largest hindwings were found primarily in upper Amazonia. Statistically, the spatial relationship between shape and elevation was particularly strong for the forewing (Table 2, Supplementary Figure S7, Supplementary Table S3). The association between wing size and elevation, however, is probably largely explained by shared ancestry since PGLS analyses revealed no correlation between wing size and species mean elevation when phylogenetic signal is accounted for (Supplementary Table S2). On average, forewing shape and, to a lesser extent, hindwing shape varied from elongated, triangular forewings in the Andes towards rounded wings in Amazonia (Figure 4, Table 2). PGLS analyses testing for a correlation between species mean elevation and shape showed a significant relationship, after accounting for phylogenetic signal (Supplementary Table S3).

**Figure 4:**
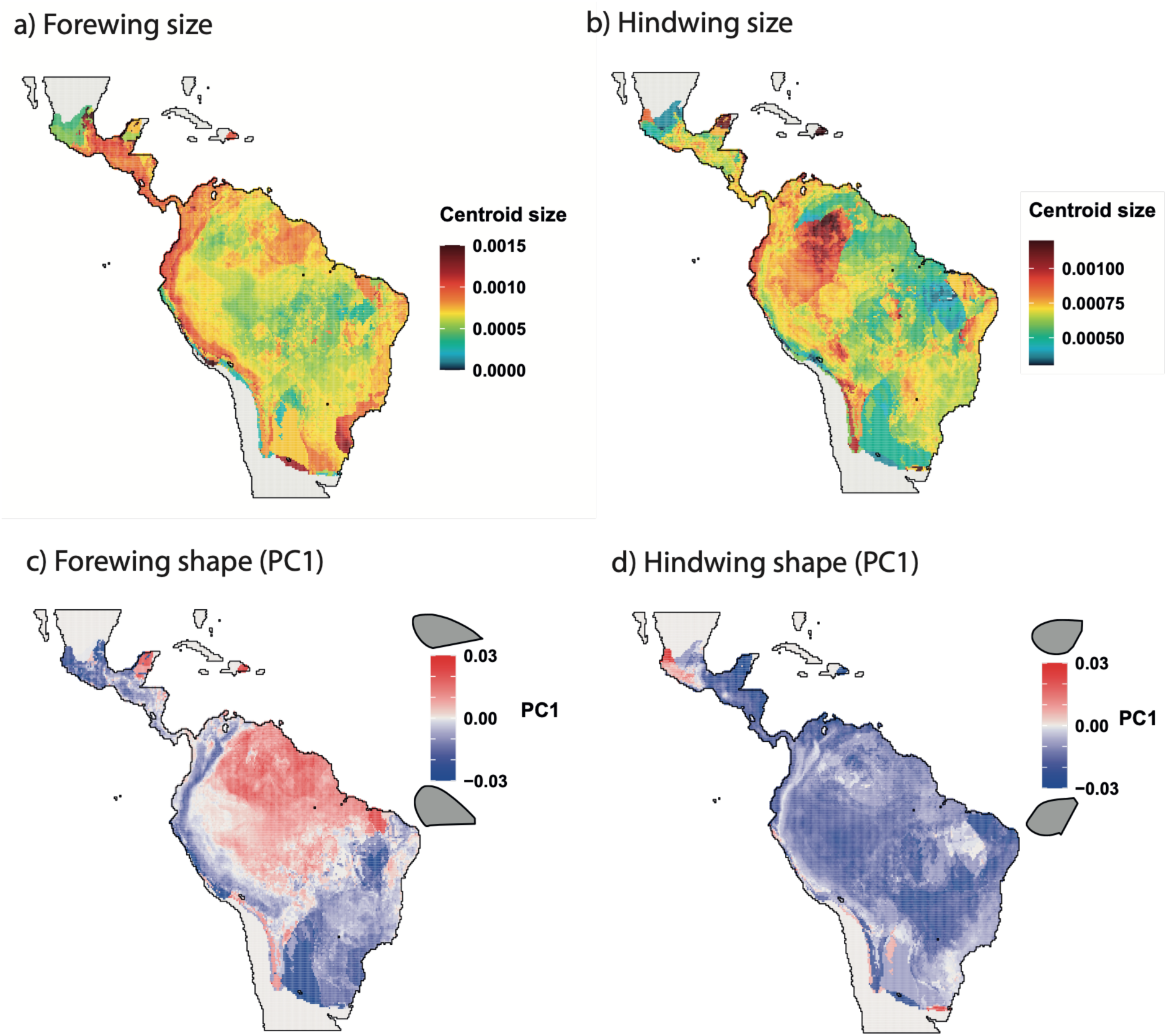
Maps of mean morphology (size and shape). Values are averaged across the local species found in each grid cell. Size and shape are derived from morphometric geometric analyses of wing shape. Size is quantified as the mean centroid size. Shape is the pPC1 axis.

**Table 2:** Spatial correlations across mean morphologies (size and shape) and elevation. Tests are Spearman’s rank correlations across grid cell values. Shape is the pPC1 axis. Elevation was aggregated from STRM remote sensing data (`elevatr` R package, Hollister, 2025). FW = forewings. HW = hindwings.

| Wing pair | Variables | $\rho$ | T-stat | df | P-value |
| --- | --- | --- | --- | --- | --- |
| FW | Elevation vs Size | 0.087 | 12.7 | 21095 | < 0.001 |
| FW | Elevation vs Shape | -0.411 | -65.5 | 21095 | < 0.001 |
| FW | Size vs Shape | -0.146 | -21.4 | 21095 | < 0.001 |
| HW | Elevation vs Size | 0.058 | 8.4 | 21095 | < 0.001 |
| HW | Elevation vs Shape | 0.127 | 18.6 | 21095 | < 0.001 |
| HW | Size vs Shape | -0.410 | -65.3 | 21095 | < 0.001 |
| HW vs. FW | Size | 0.117 | 17.1 | 21095 | < 0.001 |
| HW vs. FW | Shape | -0.238 | -35.6 | 21095 | < 0.001 |

For each wing, the geographical patterns of size and shape diversity were highly correlated (Figure 5, Table 3, Supplementary Figure S7). Forewing size and shape diversity clearly increased in the Andes and in the Atlantic Forest, whereas they were lowest in western Amazonia (Figure 5).

**Figure 5:**
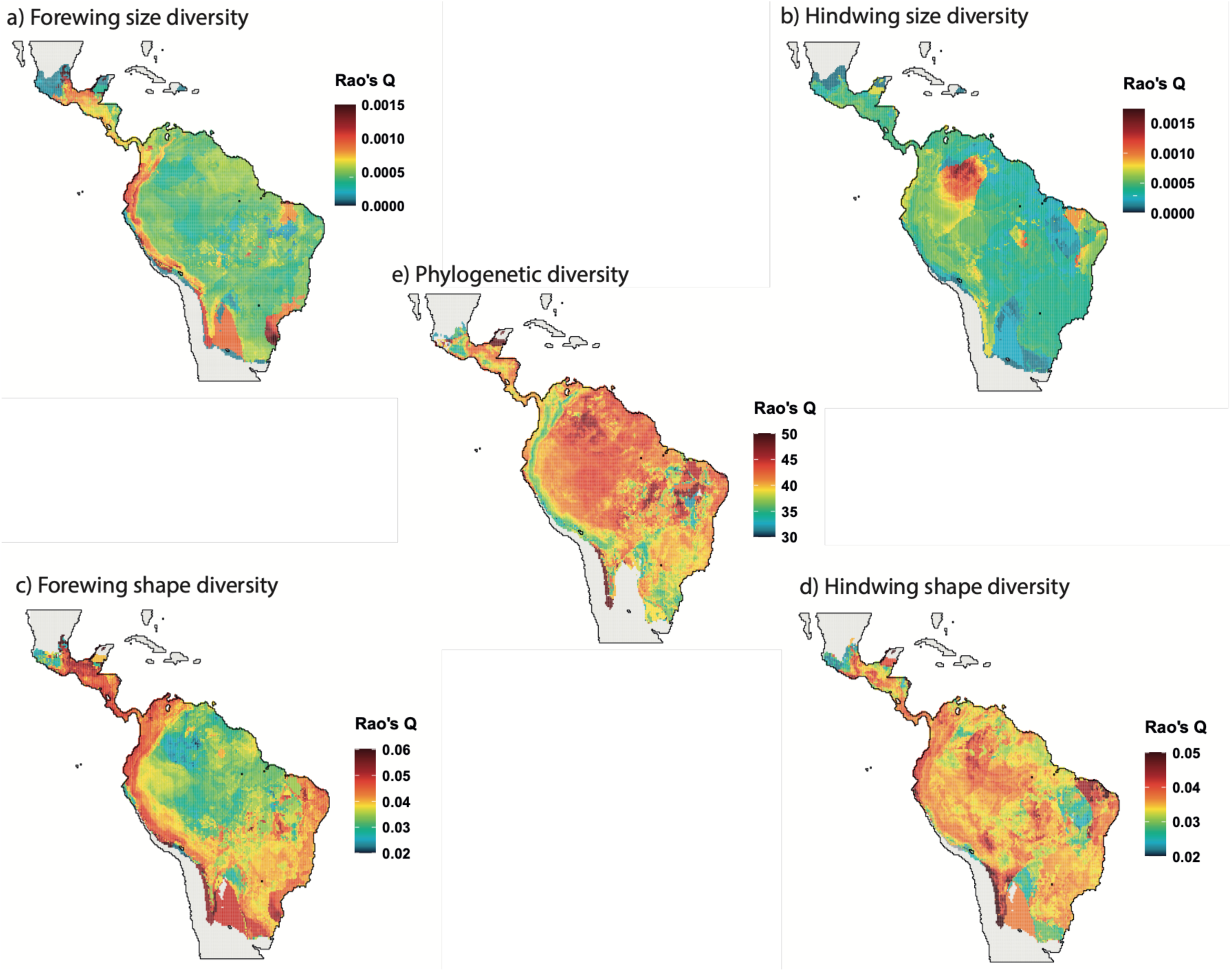
Maps of morphological (size and shape) and phylogenetic diversity estimated using Rao’s Q entropy bewteen the local species found in each grid cell. Phylogenetic diversity represents the average phylogenetic distance between local species. For size and shape diversity, this represents respectively, the average centroid size difference, and the average shape distance in the morphospace.

**Table 3:** Spatial correlations across phylogenetic and morphological diversity based on Rao’s Q entropy. Tests are Spearman’s rank correlations across grid cell values. Centroid size and pPC axes (shape) were extracted from geometric morphometrics analyses of wing shape. FW = forewings. HW = hindwings.

| Wing pair | Variables | $\rho$ | T-stat | df | P-value |
| --- | --- | --- | --- | --- | --- |
| FW | PD vs Size | -0.201 | -30.0 | 21358 | < 0.001 |
| FW | PD vs Shape | -0.331 | -51.3 | 21358 | < 0.001 |
| FW | Size vs Shape | 0.678 | 134.8 | 21358 | < 0.001 |
| HW | PD vs Size | 0.375 | 59.1 | 21358 | < 0.001 |
| HW | PD vs Shape | 0.245 | 36.9 | 21358 | < 0.001 |
| HW | Size vs Shape | 0.665 | 130.1 | 21358 | < 0.001 |
| HW vs. FW | Size | 0.001 | 0.2 | 21358 | 0.884 |
| HW vs. FW | Shape | 0.274 | 41.6 | 21358 | < 0.001 |

However, correlations between wings were weak and even non significant between forewing size diversity and hindwing size diversity. We tested whether geographical patterns of size and shape diversity were explained by phylogenetic diversity (estimated as Rao’s phylogenetic entropy, i.e. the average phylogenetic distance among species within assemblages). Forewing size and shape diversity increased where phylogenetic diversity decreased – i.e. mainly in the Andes – hindwing size and shape diversity increased with phylogenetic diversity (Table 3, Supplementary Figure S8). Of note, high scores along the margins of Ithomiini distribution result from the sensitivity of Rao’Q to the low diversity of Ithomiini inferred in these areas.

When analyzing the rate of size and shape evolution we found relatively similar spatial patterns to those identified for diversity (Figure 6, Table 4, Supplementary Figure S9). Rates of evolution of forewing size and shape were strongly correlated with one another and highly positively correlated with diversification rates (Table 4, Supplementary Figure S9). Diversification rate and phenotypic evolutionary rates clearly increased in the Andes and, by contrast, remained low in Amazonia (Figure 6). The spatial patterns of hind wing rates of evolution were more complex. The correlation between rates of evolution of size and shape was weaker than for the forewing (Table 4, Supplementary Figure S9). While the rate of shape evolution was still positively correlated between forewing and hindwing, the correlation for the rate of size evolution between wings was negative, indicating that where forewing size evolved faster, hind wing size evolution slowed down. The rate of wing shape evolution poorly correlated with species diversification rate and the rate of wing size evolution was even negatively correlated with species diversification rate (Table 4, Supplementary Figure S9).

**Figure 6:**
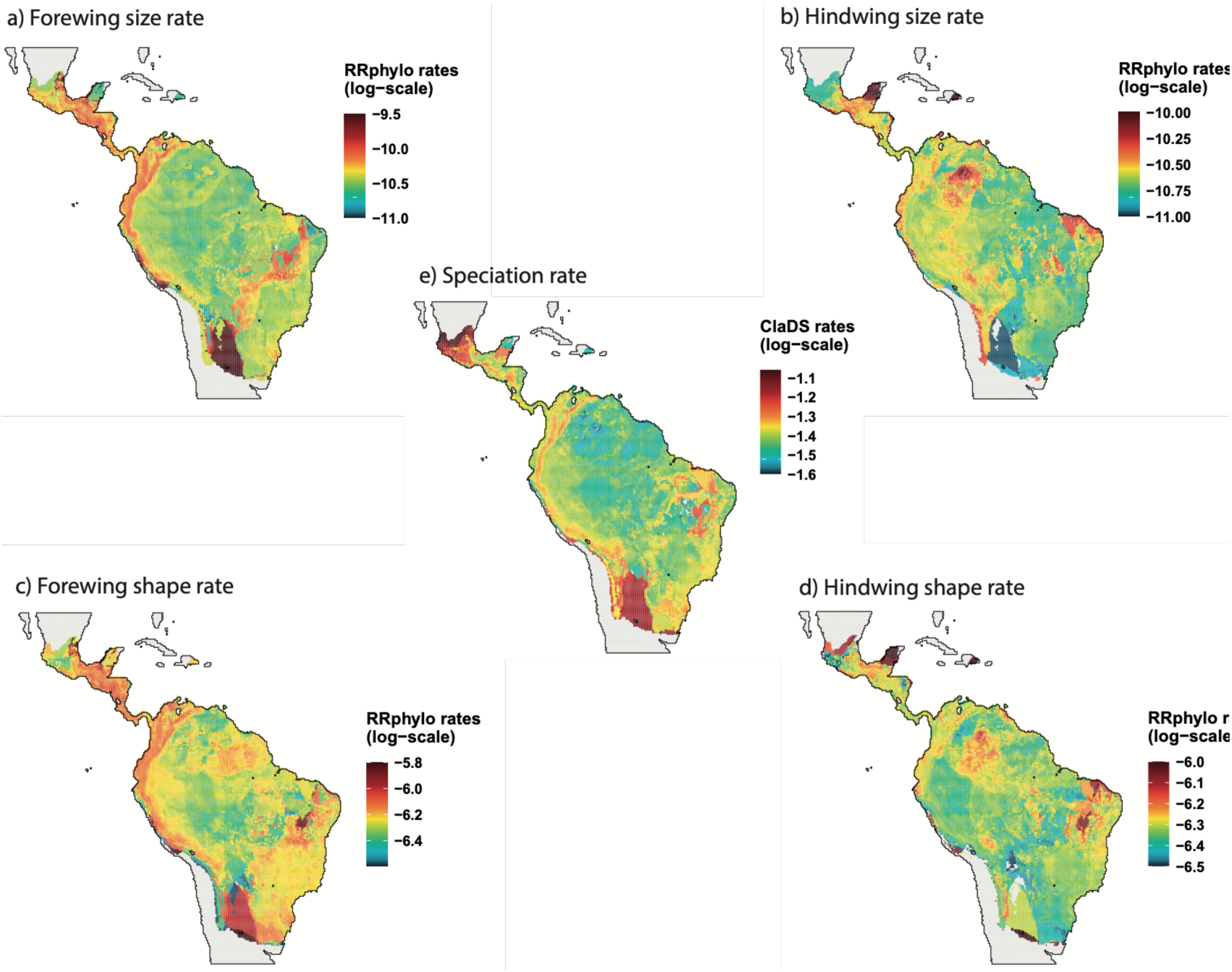
Maps of mean root-to-tip evolutionary rates: speciation, size evolution, and shape evolution. Values are averaged across the local species found in each grid cell. Diversification rates were estimated with ClaDS (Maliet et al. 2019). Size and shape evolutionary rates were estimated with RRPhylo (Castiglione et al. 2018) based on centroid size and pPC axis from geometric morphometrics analyses of wing shape. For each species, rates are average along all branches connecting the root to the species tip.

**Table 4:** Spatial correlations across root-to-tip evolutionary rates for diversification, size evolution, and shape evolution. Tests are Spearman’s rank correlations across grid cell values. FW = forewings. HW = hindwings.

| Wing pair | Rates | $\rho$ | T-stat | df | P-value |
| --- | --- | --- | --- | --- | --- |
| FW | Diversification vs Size | 0.672 | 132.7 | 21383 | < 0.001 |
| FW | Diversification vs Shape | 0.506 | 85.8 | 21383 | < 0.001 |
| FW | Size vs Shape | 0.684 | 137.1 | 21383 | < 0.001 |
| HW | Diversification vs Size | -0.251 | -37.9 | 21383 | < 0.001 |
| HW | Diversification vs Shape | 0.074 | 10.9 | 21383 | < 0.001 |
| HW | Size vs Shape | 0.373 | 58.8 | 21383 | < 0.001 |
| HW vs. FW | Size | -0.169 | -25.1 | 21383 | < 0.001 |
| HW vs. FW | Shape | 0.318 | 49.1 | 21383 | < 0.001 |

## Discussion

Our analyses reveal that wing evolution in Ithomiini is characterized by partial and context-dependent coupling among diversification rates, aspects of phenotypic evolution, and geography. Wing morphology is strongly structured by phylogeny, but the strength and consequences of this relationship differ between trait dimensions (size and shape) and between forewings and hindwings. Through time, evolutionary rate shifts are largely trait- and wing-specific, with only limited overlap between forewing and hindwing dynamics. Diversification rates are not generally associated with wing morphological evolution, although there is evidence for an association with forewing shape, whose evolutionary rates are positively associated with diversification, particularly the Andes. By contrast, hindwing morphology shows more complex and partly opposing spatial dynamics, suggesting that forewings and hindwings have followed distinct macroevolutionary trajectories.

Together, these results indicate that phenotypic diversification in Ithomiini is not uniformly associated with species diversification, but rather depends on the trait, type of wing, and geographic context considered.

### Forewings and hindwings show contrasting macroevolutionary trajectories

Our results reveal very limited coupling between forewing and hindwing evolution. We find a single shared rate shift in the evolution of size at the root of the genus *Oleria* and the rate of wing shape evolution increases for both wings in the subtribe Dircennina. The hindwing is generally much more phylogenetically conserved – phylogenetic signal for both size and shape is stronger – and, geographically, hindwing diversity appears to be determined by phylogenetic diversity. The evolution of the forewing instead appears more labile and spatially associated with diversification, especially in the Andes.

These results contrast with the few comparative studies on macroevolutionary dynamics of butterfly wing evolution. In *Papilio* swallowtails, decoupling between wings was strong, but forewing morphology was inferred to be more constrained and hindwing morphology more labile (Owen et al. 2020). In *Papilio*, hindwing shape evolves substantially faster than forewing shape, and much of this variation is associated with the presence and morphology of tails, which are thought to enhance escape from predators (Barber et al. 2015, Chotard et al. 2022, Puissant et al. 2026). Anti-predator defenses of Ithomiini differ fundamentally from those of many swallowtails, offering a possible explanation for these contrasting patterns. While *Papilio* often harbor traits such as tails, wing shape, and evasive flight that reduce predation risk, Ithomiini are chemically defended and participate extensively in Müllerian mimicry. Strong predator-mediated selection in Ithomiini may therefore act more on conspicuous wing traits involved in warning signal display and mimicry, potentially reducing the role of hindwing shape in escape-related performance. Under such a scenario, forewing morphology may be more responsive to the combined effects of mimicry, habitat structure, and Andean diversification (see discussion below), whereas hindwing morphology may retain stronger phylogenetic structure. Interestingly, Owen et al. 2020 reported that, within *Papilio*, species involved in (Batesian) mimicry are also associated with a strong reduction of tails.

A third pattern of wing evolution was reported for *Morpho* butterflies, where the two wings show strong integration, partly driven by microhabitat use (Chazot et al. 2016), and coupled dynamics of evolution through time (Chazot et al. 2021). The only signal of decoupling detected in *Morpho* butterflies appeared to be driven by sexual selection (Chazot et al. 2016). Overall, these results suggest that the relative evolutionary roles of forewings and hindwings are not fixed across butterfly radiations, but strongly depend on lineage-specific ecological and selective contexts, which in turn, enable decoupled dynamics of evolution in space and time.

### Dynamics of wing evolution and diversification are overall decoupled

Our results suggest that, in Ithomiini, wing size is more evolutionarily labile than wing shape, while shape is more strongly structured by shared ancestry or long-term constraints. There are very few points for comparison for this pattern among insects. Pie & Tschá (2013) also found that body size evolved faster than body shape in ants, but Owen et al.’s (2020) macrevolutionary study of *Papilio* swallowtail wings found that wing shape evolved faster than wing size.

The distinction between the traits of size and shape is consistent with earlier morphometric work on Ithomiini and Heliconini. Strauss (1990) showed that wing size and wing shape do not vary equivalently: forewing and hindwing areas scale isometrically, whereas wing shape is strongly allometric and differs between small and large species. Thus, even when forewings and hindwings are coupled in overall size scaling, shape variation can follow more complex and partly independent trajectories. Our results extend this static morphometric pattern to macroevolutionary dynamics: forewing and hindwing evolution shows limited covariation in evolutionary rates, at least when the geographical context is ignored.

Despite this heterogeneity in wing evolution, we also find little evidence that wing morphology is broadly coupled to speciation dynamics through time. The Cor-STRATES framework did not identify any correlation between evolutionary rates of wing size (or extant wing size) with species diversification, and hindwing shape evolutionary rates are also unrelated to diversification.

This pattern depicts a broader macroevolutionary picture in which lineage diversification and phenotypic evolution are often only partially coupled. Some comparative studies have found positive associations between morphological evolutionary rates and species richness or speciation rates, as in salamanders and ray-finned fishes, whereas others show that diversification, niche evolution, and phenotypic evolution can be temporally lagged or distributed along a continuum of coupling and decoupling across clades (Hähn et al. 2025, Morinaga & Bergmann 2023, Chazot et al. 2021, Rabosky 2013, Rabosky & Adams 2012, Folk et al. 2019, Graham & Fine 2008). In Ithomiini, the dominant pattern is therefore not a simple tree-wide relationship between speciation and wing evolution, but rather a more selective association involving particular traits, wing types, and geographic contexts.

### The Andes concentrate forewing morphological diversity and evolutionary rates

We found correlated dynamics of evolution associated specifically with the forewing and when taking into account the geographical context. The evolutionary rate dynamics of the forewing shape correlate with the rate of evolution of forewing size and with speciation rate across the tree. Our results point at a pivotal role played by the Andes as a driving force behind such correlations. We find high diversity of forewing size and shape and high evolutionary rates in the Andes for both forewing size and shape and high diversification rates. This occurs despite relatively low phylogenetic diversity, suggesting that Andean clades are phylogenetically clustered but morphologically diverse.

The evolutionary rate shift we found at the root of Godyridina and Dircennina sheds some light on this geographic signal in forewing shape evolution. These two subtribes are the most species-rich within Ithomiini and, for both of them, the Andes represent a major center of diversification (Chazot et al. 2019, Chazot et al. 2016, Lisa De-Silva et al. 2017).

There are examples of adaptive evolution of wings to elevation in Lepidoptera, such as in *Heliconius* butterflies (Montejo-Kovacevich et al. 2021, Montejo-Kovacevich et al. 2019) or *Tecia* moths (Hernández-L. et al 2010), possibly linked to flight performances under different air pressure conditions. However, why Andean diversification is associated with an increasing rate of wing evolution and the forewing specifically (and not hindwing) is unclear. One plausible hypothesis is that the Andean slopes represent a strong environmental gradient with rapid turnover of climate, vegetation, habitats and predators. Habitat and microhabitat generate strong selection on wing morphology in butterflies, with different conditions selecting, for example, for increased manoeuvrability, gliding or evasive capacity (Chazot et al. 2016; Le Roy et al., 2021). Repeated evolution across altitudinal bands and habitats during diversification may have generated fast morphological evolution.

Ithomiini can be found up to 3000 m in the Andes, where air density is significantly lower than at lower altitudes. These conditions have strong effects on the aerodynamic forces applying to butterfly wings and therefore perhaps drive the evolution of adaptations that generate lift more efficiently (Kang et al. 2023). The Andean forewing phenotype observed in Ithomiini — larger wings with lower aspect ratio and a rounder outline — closely parallels altitude-associated wing-shape evolution in *Heliconius*, where high-elevation species and populations also tend to have rounder wings (Montejo-Kovacevich et al. 2019, Montejo-Kovacevich et al. 2021). Such wings may help compensate for reduced air density by increasing wing area and lowering wing loading, while lower aspect ratio may favour manoeuvrability and force production at lower flight speeds in montane forest habitats (Dudley 2000, Betts & Wootton 1988). In Lepidoptera, forewings appear to provide the main aerodynamic force required for basic flight, whereas hindwings contribute disproportionately to manoeuvrability and evasive performance (Jantzen & Eisner 2008). This functional asymmetry may help explain why forewings mainly show adaptive patterns in relation to altitude. Direct comparisons of flight behaviour and wing kinematics between lowland and high-elevation Ithomiini would be needed to determine whether these morphological differences translate into functional flight adaptations. In particular, measuring wingbeat frequency, stroke amplitude, flight speed, manoeuvrability, and forewing–hindwing coordination across elevations could help connect the Andean forewing phenotype to its aerodynamic consequences.

## Conclusion

Our results highlight the fact that macroevolutionary coupling of diversification dynamics (of phenotypes and species) is not a general property of diversification, but a context-dependent relationship that depends on phenotype, phylogeny, and geography. In Ithomiini, species diversification is not broadly associated with wing evolution, yet this apparent decoupling masks a more specific pattern: forewing size and shape, but not hindwing, are linked to diversification only when considering the geographic context. This finding underscores the importance of decomposing complex phenotypes into biologically meaningful components. Treating wings as a single structure obscures the contrasting dynamics of forewings and hindwings, just as ignoring geography obscures the roles of montane regions such as the Andes in concentrating forewing morphological diversity and evolutionary rates. Ithomiini therefore reveal that diversification and phenotypic evolution can be coupled only in particular anatomical and spatial contexts, suggesting that the predictability of macroevolutionary dynamics depends on the scale and component of biodiversity considered.

## Notes

### Competing Interest Statement

The authors have declared no competing interest.

