## Supplementary for "Decoupled wing evolution reveals trait-specific links between morphology, geography, and diversification in Ithomiini butterflies"

### SUPPLEMENTARY INFORMATION

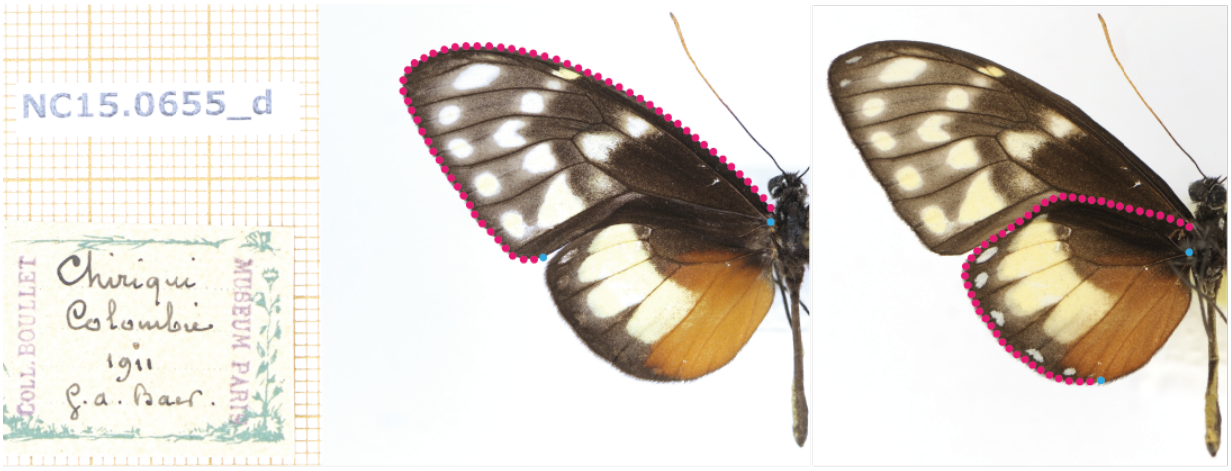


**Supplementary Figure S1**. Landmarks (blue) and sliding landmarks (red) placed for this study.

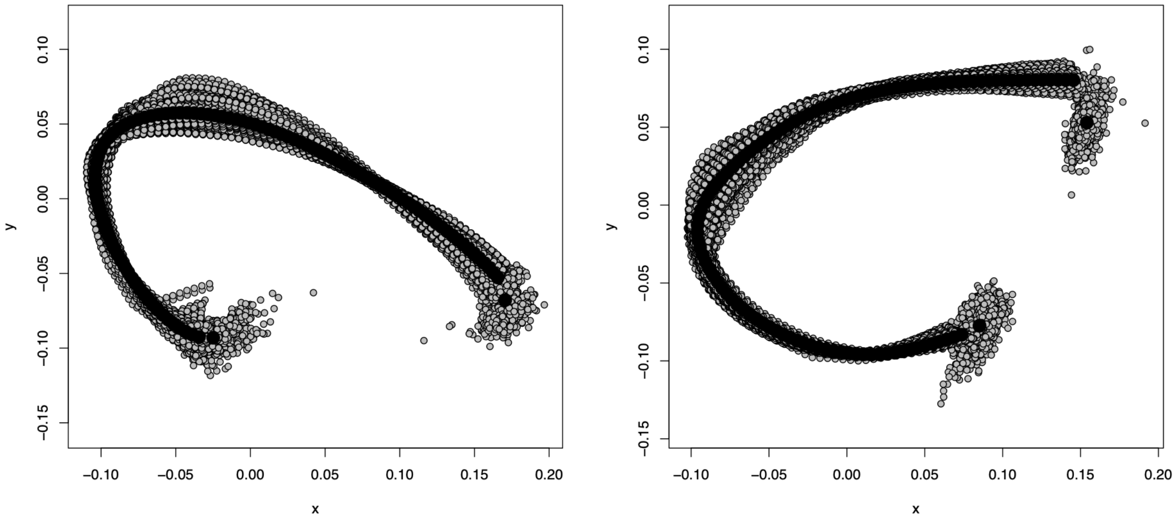


**Supplementary Figure S2**. Procrustes analysis of the forewings (left) and the hind wings (right)

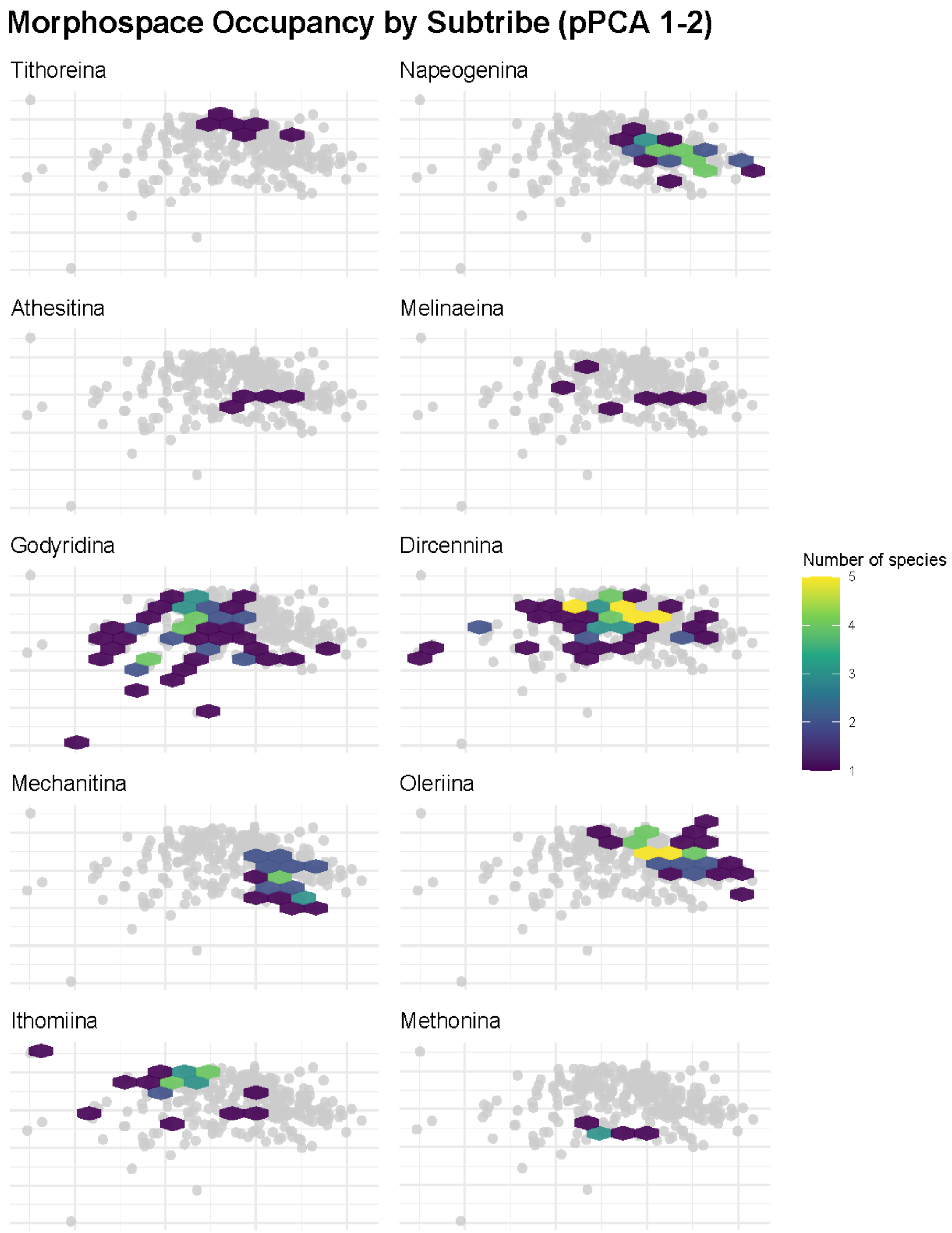


**Supplementary Figure S3**. Number of species occupying the morphospace by subtribe. Forewing phylogenetic PCA axes 1, 2.

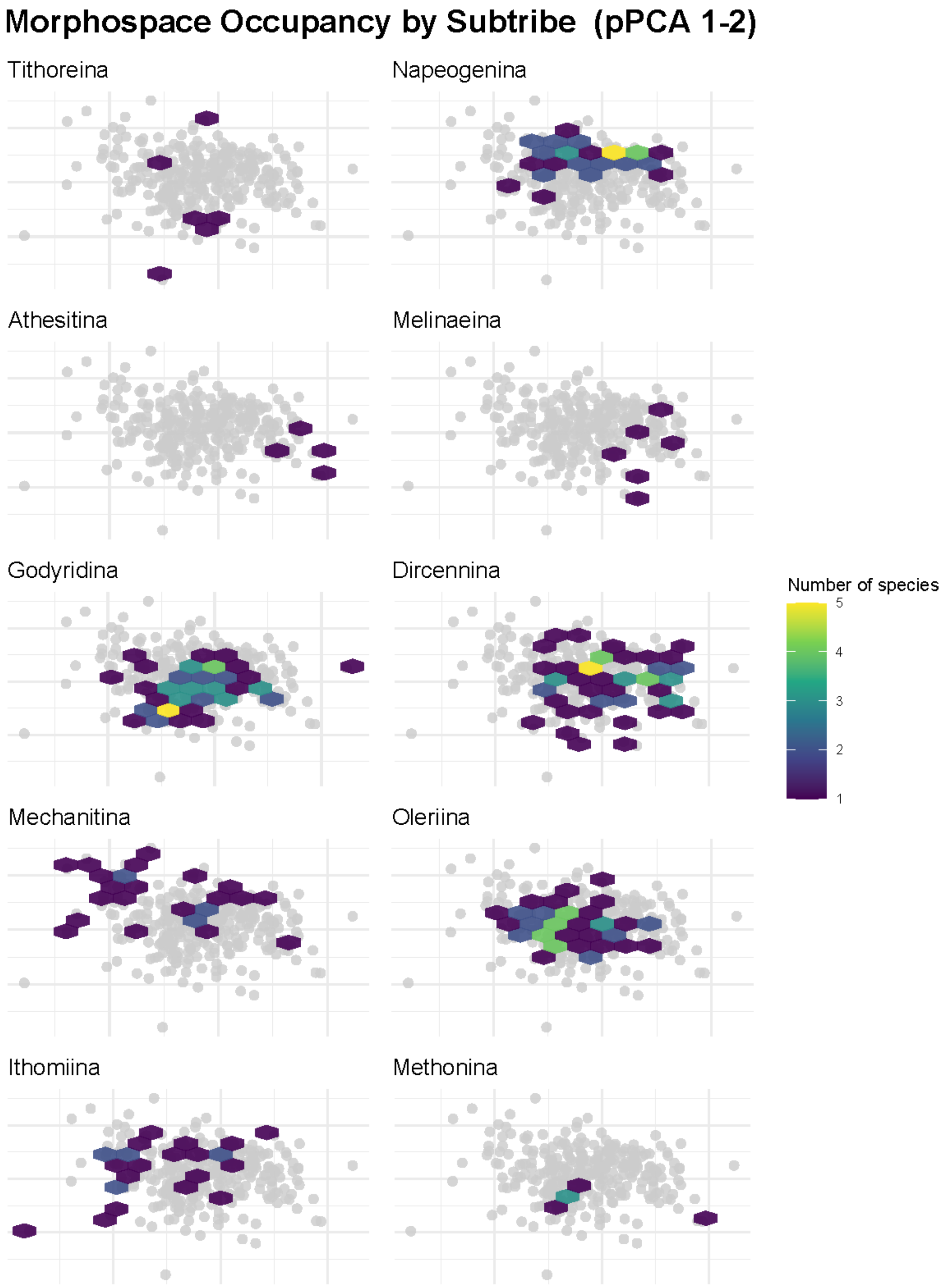


**Supplementary Figure S4**. Number of species occupying the morphospace by subtribe. Hindwing phylogenetic PCA axes 1, 2.

**Supplementary Figure S5**. Forewing (FW) and hindwing (HW) mean centroid size for each species in the tree (next page).

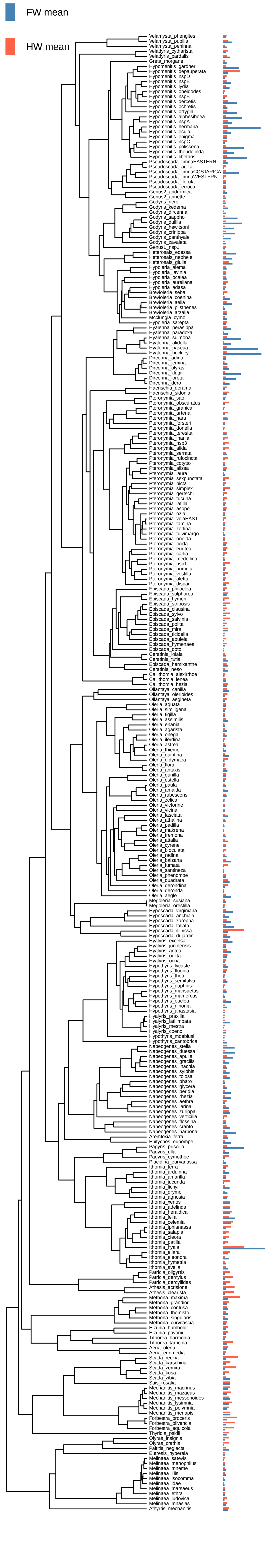

FW mean
HW mean
Velamysta_phengites
Velamysta_pupilla
Velamysta_peninna
Veladyris_cytharista
Veladyris_pardalis
Greta_morgane
Hypomenitis_gardneri
Hypomenitis_depauperata
Hypomenitis_nspD
Hypomenitis_nspE
Hypomenitis_lydia
Hypomenitis_oneidodes
Hypomenitis_nspB
Hypomenitis_dercetis
Hypomenitis_ochretis
Hypomenitis_ortygia
Hypomenitis_alphesiboea
Hypomenitis_nspA
Hypomenitis_hermana
Hypomenitis_esula
Hypomenitis_enigma
Hypomenitis_nspC
Hypomenitis_polissena
Hypomenitis_theudelinda
Hypomenitis_libethris
Pseudoscada_timnaEASTERN
Pseudoscada_acilla
Pseudoscada_timnaCOSTARICA
Pseudoscada_timnaWESTERN
Pseudoscada_florula
Pseudoscada_erruca
Genus2_andromica
Genus2_annette
Godyris_nero
Godyris_kedema
Godyris_dircenna
Godyris_sappho
Godyris_duillia
Godyris_hewitsoni
Godyris_crinippa
Godyris_panthyale
Godyris_zavaleta
Genus1_nsp1
Heterosais_edessa
Heterosais_nephele
Heterosais_giulia
Hypoleria_alema
Hypoleria_lavinia
Hypoleria_ocalea
Hypoleria_aureliana
Hypoleria_adasa
Brevioleria_seba
Brevioleria_coenina
Brevioleria_aelia
Brevioleria_plisthenes
Brevioleria_arzalia
Mcclungia_cymo
Hypoleria_sarepta
Hyalenna_perasippa
Hyalenna_paradoxa
Hyalenna_sulmona
Hyalenna_alidella
Hyalenna_pascua
Hyalenna_buckleyi
Dircenna_adina
Dircenna_jemina
Dircenna_olyras
Dircenna_klugii
Dircenna_loreta
Dircenna_dero
Haenschia_derama
Haenschia_sidonia
Pteronymia_sao
Pteronymia_obscuratus
Pteronymia_granica
Pteronymia_artena
Pteronymia_hara
Pteronymia_forsteri
Pteronymia_donella
Pteronymia_teresita
Pteronymia_inania
Pteronymia_nsp3
Pteronymia_alida
Pteronymia_serrata
Pteronymia_rufocincta
Pteronymia_cotytto
Pteronymia_alissa
Pteronymia_laura
Pteronymia_sexpunctata
Pteronymia_picta
Pteronymia_simplex
Pteronymia_gertschi
Pteronymia_tucuna
Pteronymia_latilla
Pteronymia_asopo
Pteronymia_ozia
Pteronymia_veiaEAST
Pteronymia_tamina
Pteronymia_zerlina
Pteronymia_fulvimargo
Pteronymia_oneida
Pteronymia_ticida
Pteronymia_euritea
Pteronymia_carlia
Pteronymia_medellina
Pteronymia_nsp1
Pteronymia_primula
Pteronymia_vestilla
Pteronymia_aletta
Pteronymia_dispar
Episcada_philoclea
Episcada_sulphurea
Episcada_hymen
Episcada_striposis
Episcada_clausina
Episcada_sylvo
Episcada_salvinia
Episcada_polita
Episcada_mira
Episcada_ticidella
Episcada_apuleia
Episcada_hymenaea
Episcada_doto
Ceratinia_iolaia
Ceratinia_tutia
Episcada_hemixanthe
Ceratinia_neso
Callithomia_alexirrhoe
Callithomia_lenea
Callithomia_hezia
Ollantaya_canilla
Ollantaya_olerioides
Ollantaya_aegineta
Oleria_aquata
Oleria_similigena
Oleria_tigilla
Oleria_assimilis
Oleria_enania
Oleria_agarista
Oleria_onega
Oleria_ilerdina
Oleria_astrea
Oleria_thiemei
Oleria_quintina
Oleria_didymaea
Oleria_flora
Oleria_antaxis
Oleria_gunilla
Oleria_estella
Oleria_paula
Oleria_amalda
Oleria_rubescens
Oleria_zelica
Oleria_victorine
Oleria_vicina
Oleria_fasciata
Oleria_athalina
Oleria_padilla
Oleria_makrena
Oleria_tremona
Oleria_attalia
Oleria_cyrene
Oleria_bioculata
Oleria_radina
Oleria_baizana
Oleria_fumata
Oleria_santineza
Oleria_phenomoe
Oleria_quadrata
Oleria_derondina
Oleria_deronda
Oleria_aegle
Megoleria_susiana
Megoleria_orestilla
Hyposcada_virginiana
Hyposcada_anchiala
Hyposcada_zarepha
Hyposcada_taliata
Hyposcada_illinissa
Hyposcada_dujardini
Hyalyris_excelsa
Hyalyris_juninensis
Hyalyris_antea
Hyalyris_oulita
Hyalyris_ocna
Hypothyris_lycaste
Hypothyris_fluonia
Hypothyris_thea
Hypothyris_semifulva
Hypothyris_daphnis
Hypothyris_mansuetus
Hypothyris_mamercus
Hypothyris_euclea
Hypothyris_ninonia
Hypothyris_anastasia
Hyalyris_praxilla
Hyalyris_latilimbata
Hyalyris_mestra
Hyalyris_coeno
Hypothyris_moebiusi
Hypothyris_cantobrica
Napeogenes_stella
Napeogenes_duessa
Napeogenes_apulia
Napeogenes_gracilis
Napeogenes_inachia
Napeogenes_sylphis
Napeogenes_tolosa
Napeogenes_pharo
Napeogenes_glycera
Napeogenes_peridia
Napeogenes_rhezia
Napeogenes_aethra
Napeogenes_larina
Napeogenes_zurippa
Napeogenes_verticilla
Napeogenes_flossina
Napeogenes_cranto
Napeogenes_harbona
Aremfoxia_ferra
Epityches_eupompe
Pagyris_priscilla
Pagyris_ulla
Pagyris_cymothoe
Placidina_euryanassa
Ithomia_terra
Ithomia_arduinna
Ithomia_amarilla
Ithomia_jucunda
Ithomia_lichyi
Ithomia_drymo
Ithomia_agnosia
Ithomia_xenos
Ithomia_adelinda
Ithomia_heraldica
Ithomia_leila
Ithomia_celemia
Ithomia_iphianassa
Ithomia_salapia
Ithomia_cleora
Ithomia_patilla
Ithomia_hyala
Ithomia_ellara
Ithomia_eleonora
Ithomia_hymettia
Ithomia_avella
Patricia_oligyrtis
Patricia_demylus
Patricia_dercyllidas
Athesis_acrisione
Athesis_clearista
Methona_maxima
Methona_grandior
Methona_confusa
Methona_themisto
Methona_singularis
Methona_curvifascia
Elzunia_humboldt
Elzunia_pavonii
Tithorea_harmonia
Tithorea_tarricina
Aeria_olena
Aeria_eurimedia
Scada_reckia
Scada_karschina
Scada_zemira
Scada_kusa
Scada_zibia
Sais_rosalia
Mechanitis_macrinus
Mechanitis_mazaeus
Mechanitis_messenoides
Mechanitis_lysimnia
Mechanitis_polymnia
Mechanitis_menapis
Forbestra_proceris
Forbestra_olivencia
Forbestra_equicola
Thyridia_psidii
Olyras_insignis
Olyras_crathis
Paititia_neglecta
Eutresis_hypereia
Melinaea_satevis
Melinaea_menophilus
Melinaea_mneme
Melinaea_lilis
Melinaea_isocomma
Melinaea_idae
Melinaea_marsaeus
Melinaea_ethra
Melinaea_ludovica
Melinaea_mnasias
Athyrtis_mechanitis

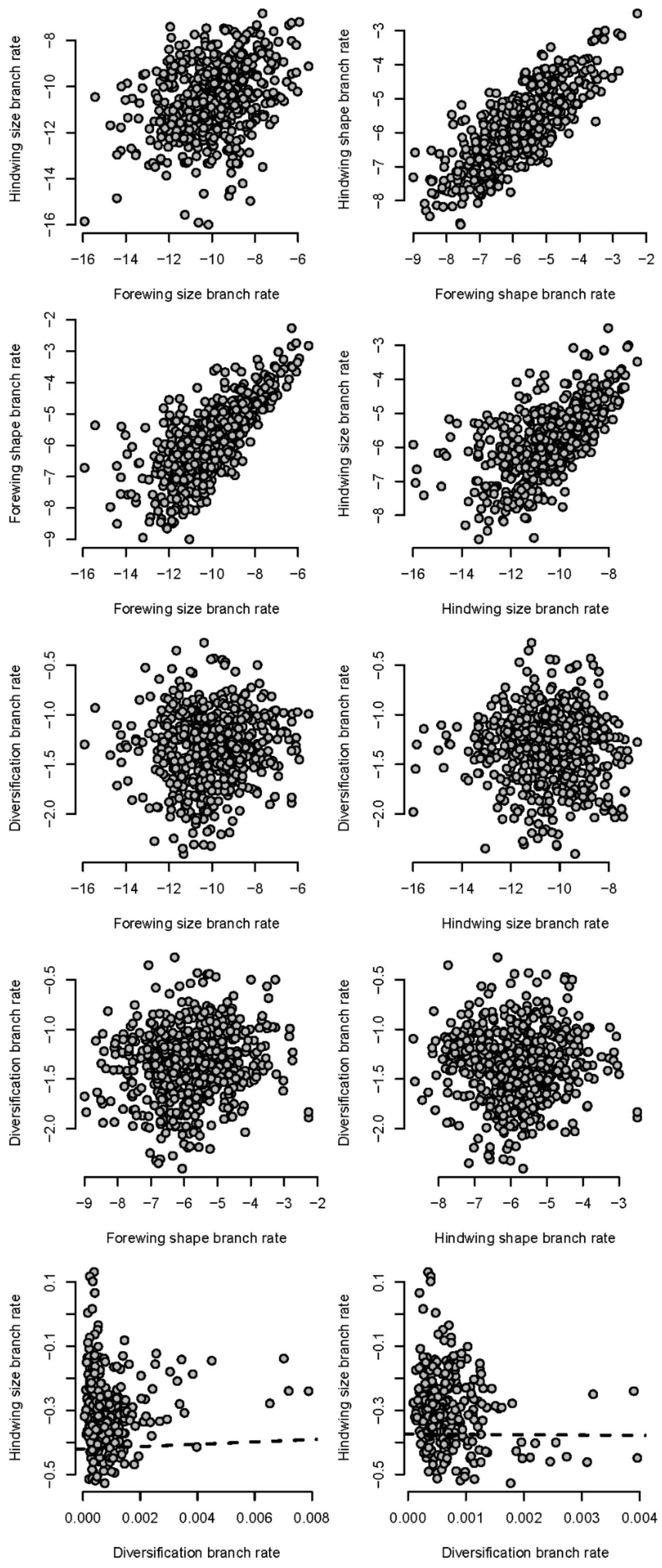


**Supplementary Figure S6**. Relationships between branch specific rate of wing size evolution, wing shape evolution, branch specific diversification rate, tip diversification rate and extant species wing size. All variables are log transformed. Phenotypic rates inferred from RRphylo are absolute values. Dashed line is the model built from PGLS analyses.

**Supplementary Table S1:** Phylogenetic generalized least-squares (PGLS) tests of the relationship between wing size and log-transformed ClaDS path diversification rate. Pagel's λ was estimated by maximum likelihood independently for each model.

| **Wing pair** | **Pagel's λ** | **Slope** | **T-stat** | **df** | **P-value** |
|---|---|---|---|---|---|
| FW | 0.449 | $6.06 \times 10^{4}$ | 0.923 | 292 | 0.357 |
| HW | 0.659 | $-5.76 \times 10^{4}$ | -1.429 | 292 | 0.154 |

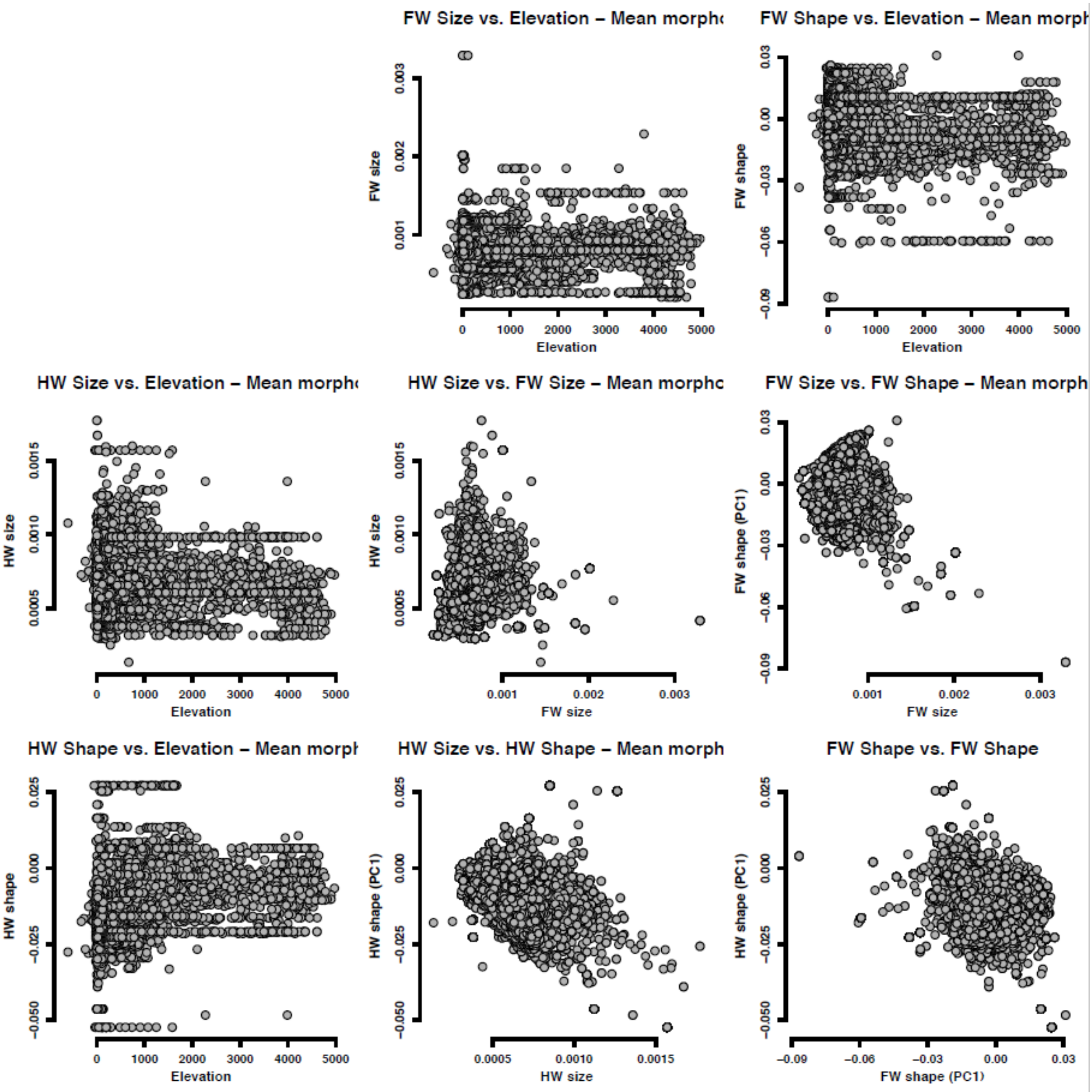


**Supplementary Figure S7:** Scatter plots of paired spatial distributions of mean morphologies (size and shape) and elevation. Data were extracted from rasters with a quarter-degree resolution covering 21,097 grid cells with at least one ithomiine species. Size and shape are derived from morphometric geometric analyses of wing shape. Size is quantified as the mean centroid size. Shape is the pPC1 axis. Elevation was aggregated from STRM remote sensing data. FW = forewings. HW = Hindwings.

**Supplementary Table S2:** Phylogenetic generalized least squares (PGLS) models testing the relationships between centroid size and altitudinal niche across species (N = 291). Models fit with R package *nlme* v3.1-168 (Pinheiro & Bates, 2025). Centroid size was log-transformed. Models included Pagel's lambda (λ) parameter to account for phylogenetic signal. Slope = Raw coefficients for the effect of elevation on size. β-coefs = Standardized coefficients for the effect of elevation on size. SE = Standard Error in parameter estimates. FW = forewings. HW = Hindwings.

| Wing pair | Pagel's λ | Slope (SE) | β-coefs (SE) | T-stat | df | P-value |
|---|---|---|---|---|---|---|
| FW | 0.438 | $9.3 \times 10^{-8}$ ($\pm$ $9.7 \times 10^{-8}$) | 0.061 (± 0.063) | 0.97 | 289 | 0.334 |
| HW | 0.681 | $1.0 \times 10^{-8}$ ($\pm$ $5.0 \times 10^{-8}$) | -0.01 (± 0.062) | -0.20 | 289 | 0.840 |

**Supplementary Table S3:** Multivariate phylogenetic generalized least squares (PGLS) models testing the relationships between shape and altitudinal niche across species (N = 291). Models fit with R package mvMORPH (Clavel et al., 2015). Shape was summarized by the first five pPC axes. Models included Pagel's lambda (λ) parameter to account for phylogenetic signal. Overall test significance was assessed through 1000 permutations comparing Wilks' statistics. Slopes = Raw coefficients for the effect of elevation on pPC shape axes. β-coefs = Standardized coefficients for the effect of elevation on pPC shape axes.

| | Forewings (FW) | Hindwings (HW) |
|---|---|---|
| **Pagel's λ** | 0.782 | 0.839 |
| **Wilks' stat** | 0.859 | 0.915 |
| **P-value** | < 0.001 | < 0.001 |
| **Effect pPC1** | | |
| Slope | $-1.8 \times 10^{-6}$ | $3.4 \times 10^{-6}$ |
| β-coef | -0.036 | 0.096 |
| **Effect pPC2** | | |
| Slope | $-2.6 \times 10^{-6}$ | $-2.4 \times 10^{-6}$ |
| β-coef | -0.15 | -0.135 |

| | | |
|---|---|---|
| **Effect pPC3** | | |
| Slope | $5.5 \times 10^{-7}$ | $2.2 \times 10^{-6}$ |
| β-coef | 0.044 | 0.156 |
| **Effect pPC4** | | |
| Slope | $-3.3 \times 10^{-6}$ | $1.4 \times 10^{-6}$ |
| β-coef | -0.324 | 0.152 |
| **Effect pPC5** | | |
| Slope | $4.7 \times 10^{-7}$ | $-5.3 \times 10^{-7}$ |
| β-coef | 0.060 | -0.066 |

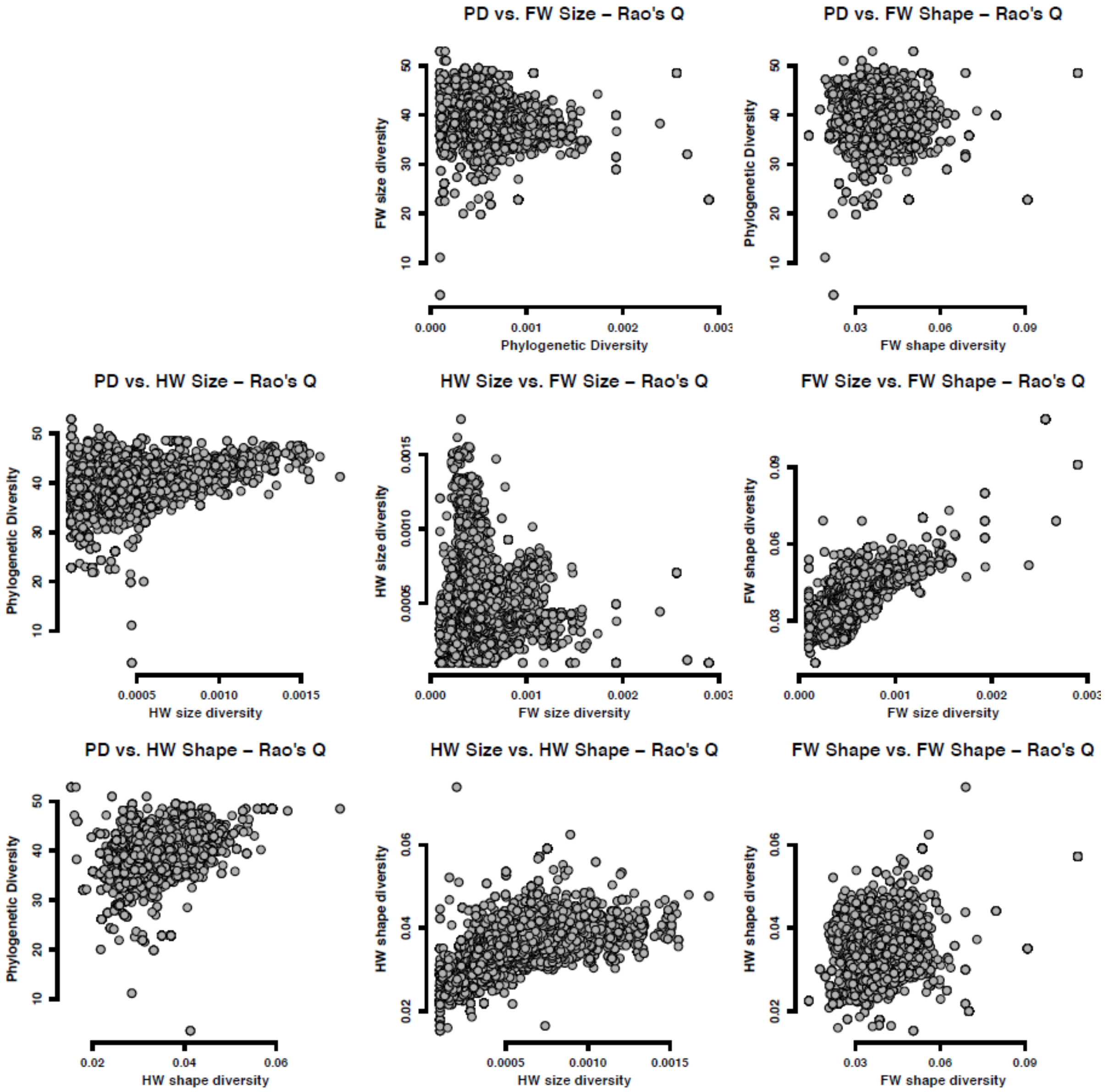


**Supplementary Figure S8:** Scatter plots of paired spatial distributions of phylogenetic and morphological diversity based on Rao's Q entropy. Data were extracted from rasters with a quarter-degree resolution covering 21,360 grid cells with at least one ithomiine species. Phylogenetic diversity represents the average phylogenetic distance between local species. For size and shape diversity, this represents respectively, the average centroid size difference, and the average shape distance in the morphospace. Centroid size and pPC axes (shape) were extracted from geometric morphometrics analyses of wing shape. FW = forewings. HW = Hindwings.

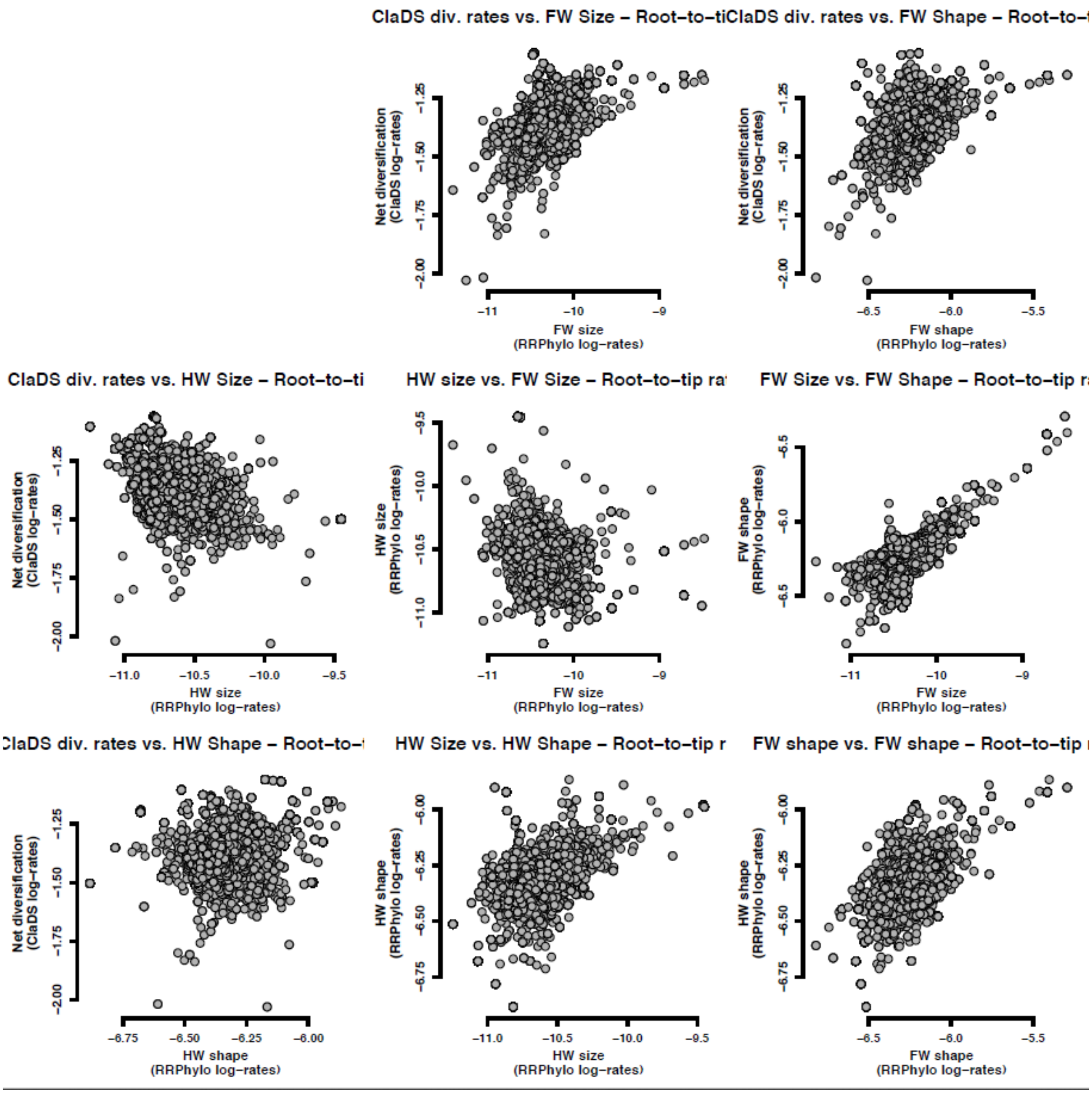


**Supplementary Figure S9:** Scatter plots of paired spatial distributions of root-to-tip evolutionary rates. Data were extracted from rasters with a quarter-degree resolution covering 21,385 grid cells with at least one ithomiine species. Diversification rates were estimated with ClaDS (Maliet et al. 2019). Size and shape evolutionary rates were estimated with RRPhylo (Castiglione et al. 2018) based on centroid size and pPC axis from geometric morphometrics analyses of wing shape. For each species, rates are average along all branches connecting the root to the species tip. FW = forewings. HW = Hindwings.
